# Adenovirus-mediated expression of SIK1 improves hepatic glucose and lipid metabolism in type 2 diabetes mellitus rats

**DOI:** 10.1101/514299

**Authors:** DaoFei Song, Lei Yin, Chang Wang, XiuYing Wen

## Abstract

**AIM:** In this study, we investigated the role and mechanism of Salt-induced kinase 1 (SIK1) in regulation of hepatic glucose and lipid metabolism in a high-fat food (HFD) and streptozocin (STZ)-induced type 2 diabetes mellitus (T2DM) rat model.

**Methods:** A diabetic rat model treated with HFD plus low-dose STZ was developed and was transduced to induce a high expression of SIK1 in vivo via a tail-vein injection of a recombinant adenoviral vector. The effects on hepatic glucogenetic and lipogenic gene expression, systemic metabolism and pathological changes were then determined.

**Results:** In T2DM rats, SIK1 expression was reduced in the liver. Overexpression of SIK1 improved hyperglycaemia, hyperlipidaemia and fatty liver, reduced the expression of cAMP-response element binding protein (CREB)-regulated transcription co-activator 2 (CRTC2), phosphoenolpyruvate carboxykinase (PEPCK), glucose-6-phosphatase (G6Pase), pS577 SIK1, sterol regulatory element binding-protein-1c (SREBP-1c) and its target genes, including acetyl-CoA carboxylase (ACC) and fatty acid synthase (FAS), and increased the expression of SIK1, pT182 SIK1 and pS171 CRTC2 in diabetic rat livers with the suppression of gluconeogenesis and lipid deposition.

**Conclusion:** SIK1 plays a crucial role in the regulation of glucose and lipid metabolism in the livers of HFD/STZ-induced T2DM rats, where it suppresses hepatic gluconeogenesis and lipogenesis by regulating the SIK1/CRTC2 and SIK1/SREBP-1c signalling pathways. Strategies to activate SIK1 kinase in liver would likely have beneficial effects in patients with T2DM and nonalcoholic fatty liver disease (NAFLD).

## Introduction

T2DM is characterized by hyperglycemia and insulin resistance (IR) and is the foremost type of diabetes around the world [1]. Diabetes complications such as hyperlipidemia and NAFLD account for an increasing proportion of annual health care costs. Tight glucose control has been associated with a reduced incidence of diabetes complications, underscoring efforts to characterize regulators that function importantly in the pathogenesis of T2DM [2].

SIK1, a serine/threonine protein kinase, belongs to the AMP-activated protein kinase (AMPK) [3]. As an energy sensor, AMPK markedly inhibits hepatic glucogenesis and lipogenesis by transcriptional control [4, 5]. In addition, Liver kinase B 1 (LKB1), a major upstream kinase of AMPK, phosphorylates SIK1 at Thr182 in the activation loop (A-loop) of the kinase domain, which is essential for switching on the SIK1 kinase activity, thus resulting in the increase of the kinase activity of SIK1 [6, 7]. Treatment with adrenocorticotropic hormone (ACTH) and the subsequent phosphorylation of the regulatory domain at Ser-577 by protein kinase A (PKA) makes SIK1 translocate to the cytoplasm and lose its repressive properties[3, 8]. Knockdown of SIK1 in mice promoted both fasting hyperglycaemia and gluconeogenic gene expression. Conversely, mice treated with adenovirus-expressed SIK1 (Ad-SIK1) exhibited fasting hypoglycaemia and reduce gluconeogenic gene expression [9]. Ad-SIK1 was also effective in reducing blood glucose levels in fasted db/db diabetic mice [9]. These observations demonstrate a key role of SIK1 on glucose metabolism in vivo.

The liver is the major organ responsible for glucose production. Hepatic glucose production mainly comes from gluconeogenesis and is critical for maintaining normoglycemia in the fasting state [10]. The cAMP response element binding protein (CREB) and its co-activator, CRTC2, play crucial roles in signal-dependent transcriptional regulation of hepatic gluconeogenesis. CREB transcriptional activity is required for fasting gluconeogenesis [11]. CRTC2 is a key regulator of fasting glucose metabolism that acts through the CREB to modulate glucose output. Phosphorylation of CRTC2 at Ser171 by AMPK results in the inhibition of the nuclear translocation of CRTC2; subsequently, the cytoplasmic localization of CRTC2 prevents its combination with CREB elements [9, 12]. Thus, gluconeogenesis is restrained. Conversely, sequestered in the cytoplasm under feeding conditions, CRTC2 is dephosphorylated and transported to the nucleus where it enhances CREB-dependent transcription in response to fasting stimuli [9]. CRTC2 has been recently found to be a substrate of SIK1 in vivo [9, 12]. SIK1 had been previously identified as a modulator of CREB-dependent transcription in adrenocortical carcinoma cells [13]. Moreover, CREB was found to occupy the SIK1 promoter in chromatin immunoprecipitation assays of primary rat hepatocytes; CRTC2 was recruited to this promoter in response to forskolin treatment [9]. The mRNA levels of CRTC2, PEPCK and G6Pase in SIK1-deficient primary rat hepatocytes were increased, while SIK1 overexpression suppressed the CRTC2 activity [9]. A recent report has shown that the selective salt-induced kinase (SIK) inhibitor HG-9-91-01 promotes dephosphorylation of CRTC2, resulting in enhanced gluconeogenic gene expression and glucose production in hepatocytes, but this effect is abolished when an HG-9-91-01-insensitive mutant SIK is introduced [14]. Interestingly, in primary rat hepatocytes, SIK1 phosphorylated CRTC2 at Ser 171 and in turn promoted its export to the cytoplasm, thereby inhibiting the expression of downstream gluconeogenic genes such as PEPCK and G6Pase [9], suggesting that regulation of CRTC2 activity by SIK1 may be crucial for inhibiting excessive hepatic glucose output. Therefore, the SIK1/CRTC2 signalling pathway will probably represent a novel strategy for suppressing hepatic gluconeogenesis and ameliorating hyperglycaemia. Although SIK1 is implicated in regulation of CRTC2 and hepatic glucose output, the glycometabolism of the kinase remains uncharacterized in the HFD/STZ-induced T2DM rat model.

In addition, the liver is also one of the major organs regulating lipid metabolism [15]. Hepatic lipogenesis contributes to accumulation of fat in the liver [16]. SREBP-1c acts as a master transcriptional regulator for the hepatic lipogenesis by activating its target genes, such as FAS and ACC. Moreover, SREBP-1c is shown to be a direct substrate for SIK1 in vitro [17]. SIK1 blocks lipogenesis by direct phosphorylation of SREBP-1c on multiple serine residues [12]. Ectopic expression of SIK1 in mouse livers reduces lipogenic gene expression and hepatic triglyceride accumulation [12]. This effect was reversed by co-expression of a phosphorylation-deficient Srebp1-c mutant [17]. A previous report has shown that lipogenic genes, such as FAS and ACC, are up-regulated by SIK1 knockdown in mouse liver, whereas overexpression of SIK1 reduces expression levels of SREBP-1c target genes, suggesting that SIK1 could regulate lipogenic gene transcript [17]. SIK1-induced phosphorylation of SREBP-1c at Ser329 is thought to be critical for the suppression of SREBP-1c transcription activity [17]. Our previous study demonstrated that overexpression of SIK1 suppressed the expression of SREBP-1c and its target genes in HepG2 cells cultured in a high glucose environment [18]. Thus, modulation of SREBP-1c activity by SIK1 would provide an attractive means for the regulation of hepatic lipogenesis. In the diabetic conditions, normal regulation of gluconeogenesis and lipogenesis is disrupted; hence the SIK1/CRTC2 and SIK1/SREBP-1c pathways may serve as therapeutic targets to modulate metabolic disorders in diabetic patients with NAFLD.

To date, the role and mechanism of SIK1 in the liver of the HFD/STZ-induced T2DM rat model remains completely unknown. Because the diabetic rat model treated with HFD plus low-dose STZ replicates the natural history and metabolic characteristics of human T2DM and develops most of the biochemical and pathological symptoms [19–24] associated with T2DM in humans, the diabetic rat model is particularly suitable for pharmaceutical research [25]. Thus, it is of interest to define the effect of SIK1 on hepatic gluconeogenesis and lipogenesis of the HFD/STZ-induced T2DM rat. In the present study, we generates a diabetic rat model treated with HFD plus low-dose STZ and focuses on the role of SIK1 in the hepatic gluconeogenic and lipogenic pathways and their effect on the resulting phenotype of lower fasting glucose levels and ameliorated fatty liver disease. Meanwhile, we use a recombinant SIK1-expressing adenovirus to obtain a high expression of SIK1 in vivo, and then assess its affect on diabetes in the HFD/STZ-induced T2DM rat model. To our knowledge, this is the first study to examine the effects of adenovirus-mediated SIK1 overexpression on hepatic glucose and lipid metabolism in the HFD/STZ-induced T2DM rats.

## Materials and Methods

### Recombinant adenovirus production

Ad-Sik1 and negative control adenovirus containing green fluorescent protein (Ad-GFP) were purchased from Gene Chem Co., Ltd. (Shanghai, China). Ad-Sik1 and Ad-GFP were obtained with a titre of 1×10^11^ plaque forming units (PFU) /ml. The recombinant adenovirus was stored at −80°C until use. Construction of both vectors was described in S1 Appendix. Sik1-overexpressing rats were established by an injection of Ad-Sik1 or Ad-GFP at an optimized dose of 5×10^9^ PFU in 50μl (diluted with physiological saline) via tail vein once a week for 8 weeks according to the manufacturer’s protocols. Meanwhile, the rats of the control and model groups received physiological saline at the same dosage by tail vein injection.

### Animal treatments

Thirty male wistar rats, three to four-weeks-old, weighing approximately 70-100 g, were supplied by BEI JING HFK BIOSCIENCE CO., LTD (Beijing, China). The protocol for using animals was approved by the research Ethics Committee of Tongji Medical College, Huazhong University of Science and Technology (Protocol Number: 822). All animals were housed with two rats per cage in an air-conditioned room (22°C ± 3°C, 50%-60% relative humidity) with a 12:12-hour light-dark cycle and were initially fed normal chow and allowed to adapt to their environment for 1 week. After acclimatization, all rats were randomly assigned to 2 groups. The control rats were fed ad libitum with a normal diet and the other rats were fed ad libitum with a HFD to induce diabetes [26]. Four weeks later, rats on HFD were injected with 36 mg/kg STZ (dissolved in citrate buffer, pH 4.5) intraperitoneally. Meanwhile, the control rats were injected with the same volume of citrate buffer. Diabetes was defined by fasting serum glucose >11.1 mmol/L 72 h after STZ injection. The diabetic rats were randomly divided into three groups: diabetes mellitus (DM) group (n=6), Ad-Sik1 group (n=8), and Ad-GFP group (n=6). The Ad-Sik1 and Ad-GFP groups received an injection of Ad-Sik1 or Ad-GFP at an optimized dose of 5×10^9^ PFU via tail vein once a week for 8 weeks. The DM and normal control groups were given an equal volume of normal saline. The normal chow (13.68%, 64.44%, and 21.88% of calories derived, respectively, from fat, carbohydrate, and protein) was provided by the Laboratory Animal Center, Huazhong University of Science and Technology (Wuhan, China). The high fat diet (rodent diet with 45% kcal fat) was purchased from WANQIANJIAXING BIOTECHNOLOGY CO., LTD (Wuhan, China). Food intake, water intake and blood glucose were monitored periodically. After an 8-week treatment, the rats were weighed and sacrificed. Fasting blood was collected from the ventral aorta and serum was separated for biochemical analysis. The liver was removed and weighed. Part of the liver was fixed in 4% paraformaldehyde and embedded in paraffin for hematoxylin and eosin (HE) staining and immunohistochemical analysis. The rest of the liver was washed with normal saline and stored at −80°C for RT-PCR and Western blot.

### Reagents

STZ was purchased from Sigma (SaintLouis, Missouri, USA). RNAiso Plus was purchased from TaKaRa (Dalian, China). SIK1 antibody was purchased from Novus Biologicals, LLC (Cat #: 82417, Littleton, USA). SREBP-1c antibody, G6Pase and CRTC2 (S171) antibody were purchased from Abcam (Cat #: ab28481, Cat #: ab83690 and Cat #: ab203187, Cambrige, UK). CRTC2 antibody, SIK1 (S577) antibody, SIK1 (T182) antibody, FAS antibody and ACC antibody were purchased from Proteintech Group, Inc. (Cat #:12497-1-AP, Cat #: S4530-2, Cat #: S4529-2, Cat #:10624-2-AP and Cat #:21923-1-AP, Rosemont, USA). PEPCK antibody and β-actin antibody were purchased from Cell Signaling Technology, Inc. (Cat #: 12940 and Cat #: 4967, Danvers, Massachusetts, USA). Goldview DNA dye and DNA Marker I were purchased from TIANGEN BIOTECH (BEIJING) CO., LTD. Horseradish peroxidase-conjugated goat anti-rabbit IgG was purchased from Bioworld Technology, Inc. (Minnesota, USA) as secondary antibody.

### Biochemical assay

Serum glucose was measured using enzymatic glucose-oxidase kits (Ruiyuan Biotechnology Co., Ltd, Ningbo, China), TG and total cholesterol (TC) were determined using enzymatic couple colorimetric kits (Huachen Biochemical Co., Ltd, Shanghai, China).

### Histological Analysis

Liver tissues were fixed in 4% paraformaldehyde for 24 h, embedded in paraffin, sectioned into 4 μm sections (Leica, Wetzlar, Germany), and stained with HE for microscopic assessment (Olympus, Tokyo, Japan). The liver cryosections were prepared for oil red O staining.

### Immunohistochemistry analysis

The liver tissues were fixed with 4% paraformaldehyde for paraffin embedding. The paraffin-embedded sections were subjected to immunohistochemical staining for SIK1, CRTC2, PEPCK, G6pase, SREBP-1c, FAS and ACC in the liver. The tissue sections were incubated with rabbit anti primary antibody (1:100). After washing with PBST, the sections were incubated with secondary antibody, and the diaminobenzidine method was used. Next, the SIK1, CRTC2, PEPCK, G6pase, SREBP-1c, FAS, ACC and Insulin protein expressions were observed under an optical microscope. All the sections were examined by light microscope. Optical density (OD) was identified as expression intensity of positive staining in the liver tissues, which was semiquantitatively analysed with Image-Pro Plus 6.0 software (Media Cybernetics, Inc., USA).

### RT-PCR

Total RNA samples were isolated from the rat liver using RNAiso Plus (D9109; TaKaRa, Dalian,China). RNA samples were converted to cDNA using a RevertAid First Strand cDNA Synthesis Kit (Thermo Fisher Scientific, USA) according to the manufacturer’s instructions. Primers were designed using the nucleotide sequence and synthesized by Sangon Biotechnology Co., Ltd. (Shanghai, China). Semi-quantitative PCR conditions were 95°C for 2 minutes, 94°C for 30 seconds, followed by 35 cycles at 55°C for 30 seconds, 72°C for 30 seconds, and 72°C for 2 minutes. PCR products were separated by 1.5% agarose gel electrophoresis and visualised under ultraviolet light using a JS-680B Bio-Imaging System camera (PEIQING Technology Limited, Shanghai, China). Relative band intensities of each sample were calculated after being normalized to the band intensity of β-actin. The sequences of the primers used in this study are listed in S1 Table.

### Western blot

SIK1, SIK1 (S577), SIK1 (T182), CRTC2, CRTC2 (S171), PEPCK, G6pase, SREBP-1c, FAS and ACC protein expression were determined by Western blotting, which was performed according to standard procedures. The protein concentration in tissue lysates was measured with a BCA protein assay kit (Boster, Wuhan, China) according to the manufacturer’s instructions. Protein lysates extracted from the rat liver tissue were electrophoresed using 8-12% sodium dodecyl sulphate-polyacrylamide gel electrophoresis (SDS-PAGE) for separation. Then, samples were transferred onto a nitrocellulose membrane. The membrane was next incubated in 5% milk in a mixture of Tris-buffered saline and Tween 20 (TBST) for 1 h at room temperature to block the membrane. The proteins were incubated with the primary antibody overnight at 4°C (SIK1, 1:1000; SIK1 (S577), 1:2000; SIK1 (T182), 1:2000; CRTC2, 1:500; CRTC2 (S171), 1:500; PEPCK, 1:1000; G6pase, 1:800;SREBP-1c, 1:500; FAS, 1:500; ACC, 1:500; β-actin, 1:1000). After washing the membrane 5 times in TBST, the membrane was incubated with secondary antibody for 30 min at room temperature. Finally, the protein was detected with electrochemiluminescence (ECL) Western blotting reagents. The optical density (OD) of protein bands was quantified using Image J 1.48 software (National Institutes of Health, USA). The results are expressed as the ratio between the OD value of a target band to the OD value of β-actin.

### Statistical analysis

All data are analysed by one-way ANOVA or two-way ANOVA using GraphPad Prism 5.0 software (GraphPad Software, San Diego, CA, USA), and the results are presented as the mean ± SD. P<0.05 was considered statistically significant. In the figures and tables, *P< 0.05; **P< 0.01; ***P< 0.001. ns, not significant.

## Results

### Effects on body weight, liver weight, FBG, TG and TC

**Figure 1.**
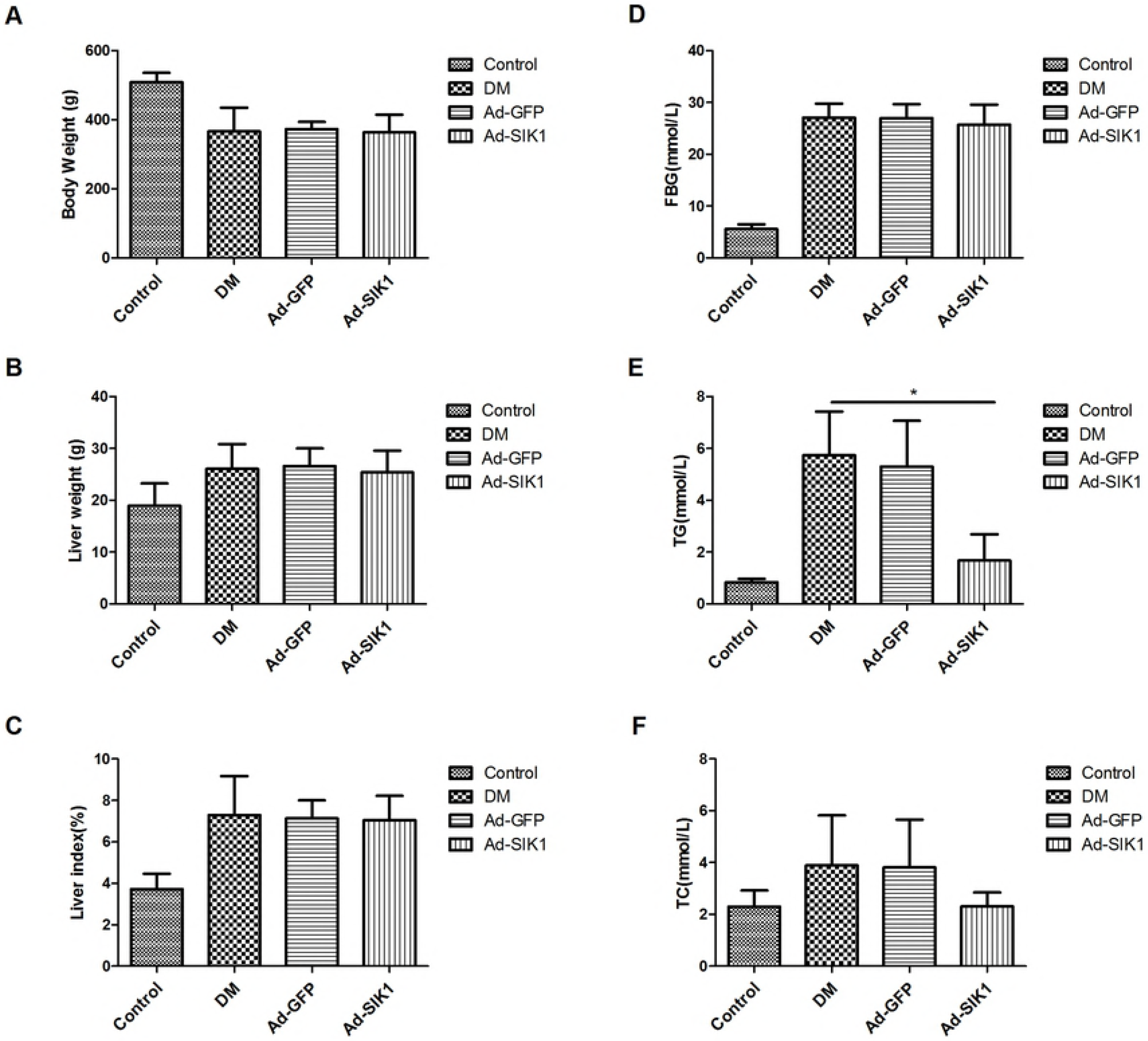
Effects on body weight, liver weight, FBG, TG and TC in HFD/STZ-induced diabetic rats. (A) Body weight; (B) Liver weight; (C) Liver index; (D) Serum glucose levels; (E) Serum TG levels; (F) Serum TC levels. The results are expressed as the mean ±SD. Significant differences are indicated as *P<0.05, **P<0.01, *** P<0.001. ns, not significant.

The HFD/STZ-induced diabetic rats showed classic diabetic symptoms of polyuria, polydipsia and weight loss. These symptoms are related to the presence of hyperglycaemia (blood glucose level fluctuation from 20.09 to 30.61 mmol/L). Neither the Ad-SIK1 group nor the Ad-GFP group showed significant differences in blood glucose or body weight. As shown in Table 1, serum glucose, TG and TC were significantly higher in the DM group compared to the control group. Intriguingly, serum TG was remarkably decreased (P<0.05), but serum TC slightly reduced in the Ad-SIK1 group compared to the DM group. Although Ad-SIK1 administration attenuated the HFD/STZ-induced increase in the serum TC level, no significant difference was observed.

### Histological examination of liver

**Fig 2.**
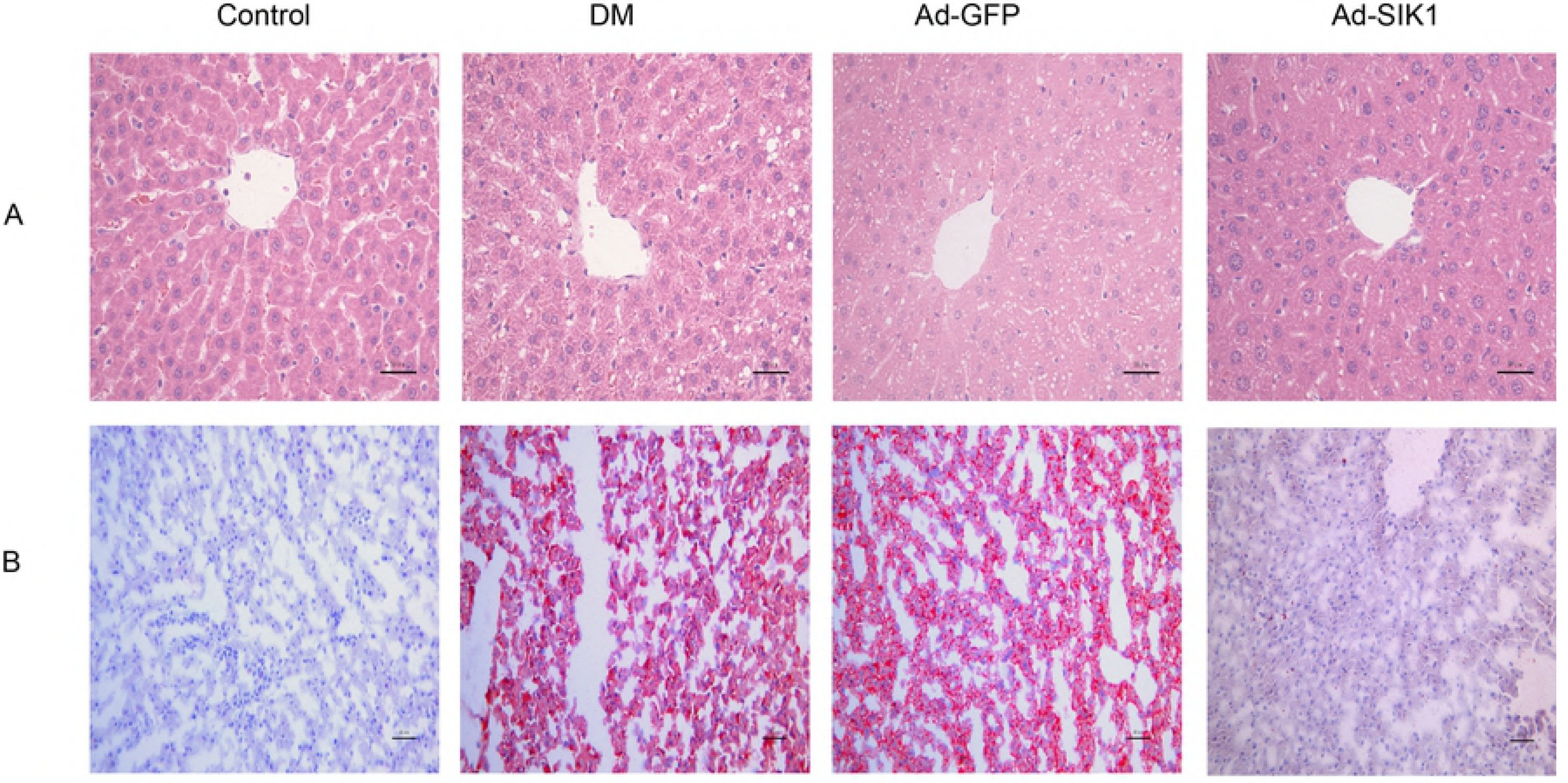
Effects on histology of liver of HFD/STZ-induced diabetic rats. (A) Liver tissue sections were stained with HE (×200); (B) oil red O to observe liver lipid content. Scale bar is 50 μm.

The typical HE and oil red O staining results obtained upon histological examination are shown in Fig 1. In consonance with the biochemical data, the staining of liver tissues with HE and oil red O revealed an accumulation of lipid droplets in the liver of the DM group, whereas lipid droplets were rare in the liver of the Ad-SIK1 group. Thus, Ad-SIK1 treatment significantly reduced fat deposition compared with the DM group, indicating that Ad-SIK1 administration could markedly improve steatosis. Therefore, these results confirmed the protective effect of SIK1 overexpression against histological changes in the liver of HFD/STZ-induced diabetic rats.

### Immunohistochemical staining of genes related to hepatic glucose and lipid metabolism

**Fig.3.**
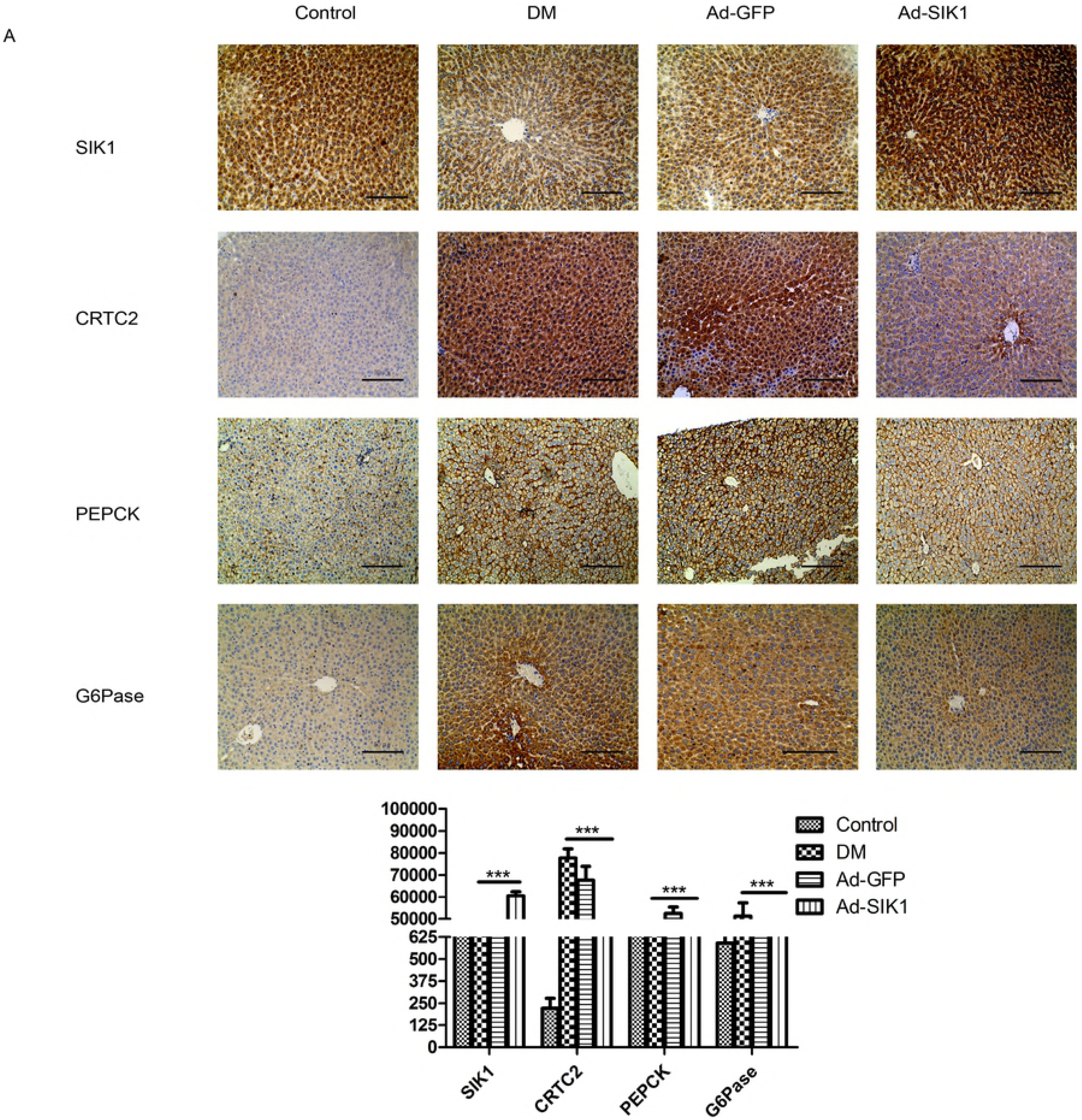
Effects on immunohistochemical staining of SIK1, CRTC2, PEPCK, G6pase, SREBP-1c, FAS and ACC in liver. (A) immunohistochemical staining of **SIK1, CRTC2, PEPCK and G6pase; (B)** immunohistochemical staining of **SREBP-1c, FAS and ACC.** Immunohistochemical staining images are 200 times larger under light microscopy Data are presented as the mean ± SD. Significant differences are indicated as * p<0.05, ** p<0.01, *** p<0.001. ns, not significant. Scale bar is 50 μm.

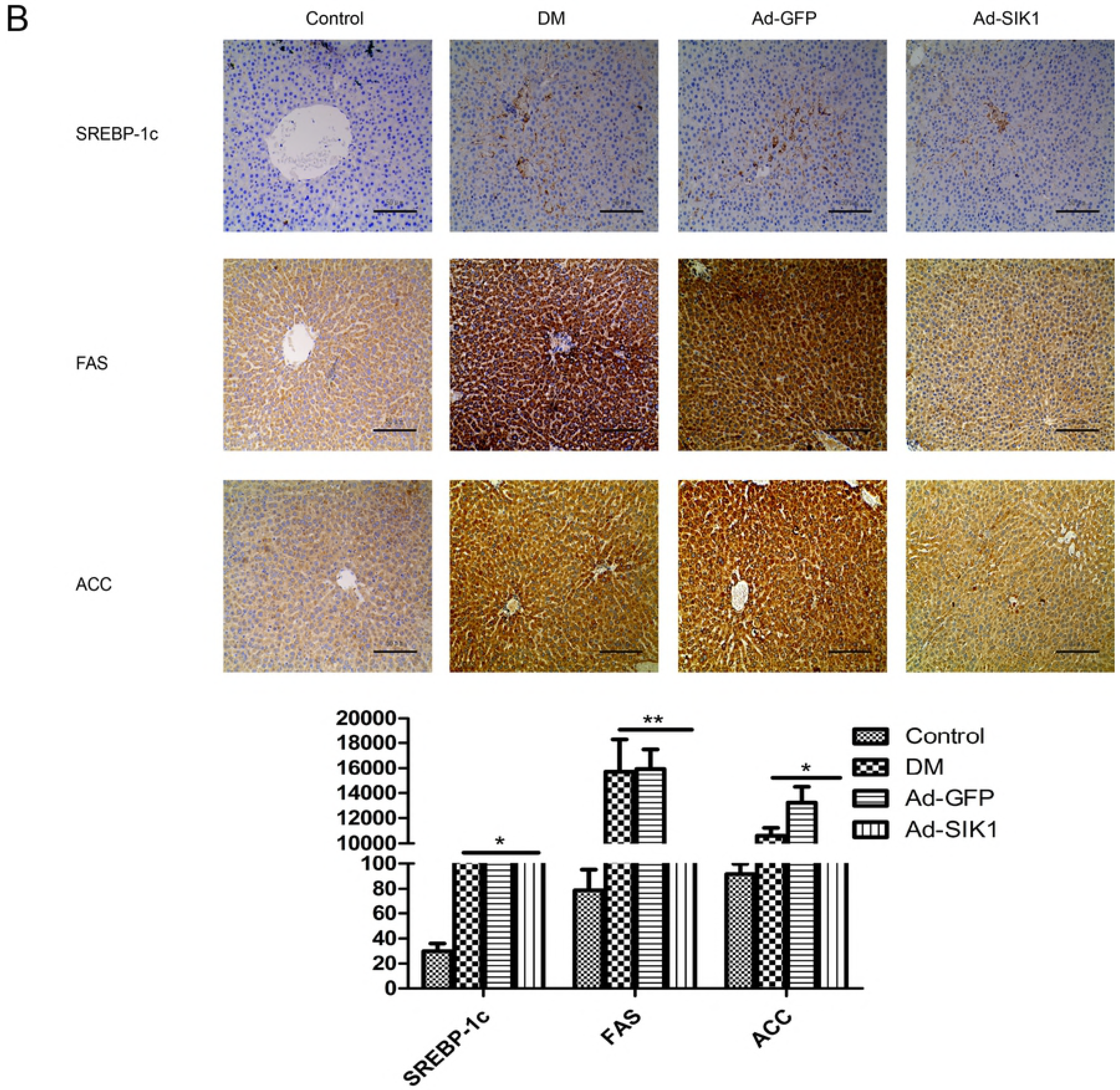

Fig 3 illustrates the immunohistochemical photomicrographs of SIK1, CRTC2, PEPCK, G6pase, SREBP-1c, FAS and ACC in the liver of rats. In the DM and Ad-GFP groups, SIK1-positive staining was much weaker than that in the Ad-SIK1 (p<0.001). Notably, SIK1 overexpression significantly increased the reduced SIK1-positive staining in the liver of diabetic rats. In contrast, CRTC2, PEPCK, G6pase, SREBP-1c, FAS and ACC stainings were much stronger in the DM and Ad-GFP groups than those in the Ad-SIK1 group (p<0.001). Obviously, Ad-SIK1 treatment inhibited this enhanced positive staining. In addition, we also verified that SIK1 overexpression inhibited CRTC2 nuclear translocation in the liver tissues *via* immunohistochemical staining. The nuclear expression of CRTC2 protein was obviously increased in the DM and Ad-GFP groups compared to the normal control group; however, the treatment with Ad-SIK1 inhibited the nuclear translocation of the CRTC2 protein.

### SIK1 overexpression results in the inhibition of the hepatic gluconeogenic program in HFD/STZ-induced diabetic rats

**Fig 4.**
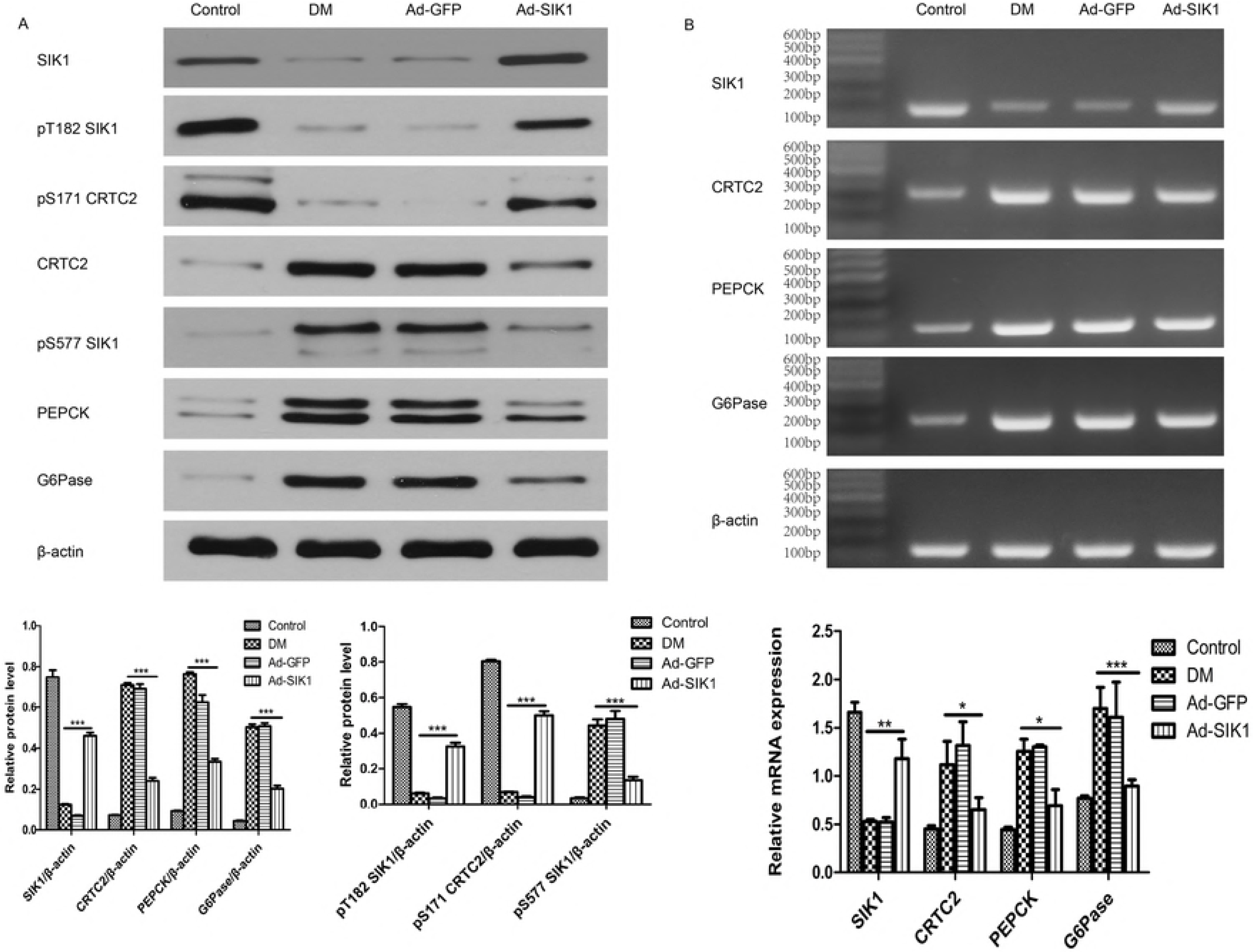
Effects on mRNA and protein expression of genes related to glucose metabolism in diabetic rats. (A) Protein levels of SIK1, CRTC2, PEPCK and G6pase in liver; (B) mRNA (relative fold change) levels of SIK1, CRTC2, PEPCK and G6pase in liver. Fold expression levels were measured relative to the expression of β-actin (internal control). Data are presented as the mean ± SD. Significant differences are indicated as * p<0.05, ** p<0.01, *** p<0.001. ns, not significant. n=6 in the control group; n=5 in the DM group; n=6 in the Ad-SIK1 group.

To determine whether SIK1 overexpression ameliorates hyperglycaemia by decreasing endogenous glucose production in the liver, we measured the mRNA and protein of SIK1, CRTC2, PEPCK and G6pase in the liver. RT-PCR analysis showed that SIK1 was significantly elevated in the Ad-SIK1 group compared to the DM group, while CRTC2, PEPCK and G6pase were significantly reduced, indicating an inhibitory effect of SIK1 in liver glucogenesis (Fig 3B). Indeed, the Western blot results showed that SIK1 was significantly decreased in the DM group compared to the control group, whereas CRTC2, PEPCK and G6pase were significantly elevated in the DM group. SIK1 overexpression significantly increased the protein level of SIK1, but decreased the protein levels of CRTC2, PEPCK and G6pase in liver compared with the DM group (Fig 3A). Meanwhile, the phosphorylation level of SIK1 at Ser577 was drastically reduced, whereas the level of pT182 SIK1 and pS171 CRTC2 was significantly increased in the Ad-SIK1 group compared with the DM and Ad-GFP groups. These results suggest that SIK1 could inhibit the hepatic gluconeogenic program in HFD/STZ-induced diabetic rats by regulating the SIK1/CRTC2 signalling pathway.

### SIK1 inhibits the hepatic lipogenic program in HFD/STZ-induced diabetic rats

**Fig 5.**
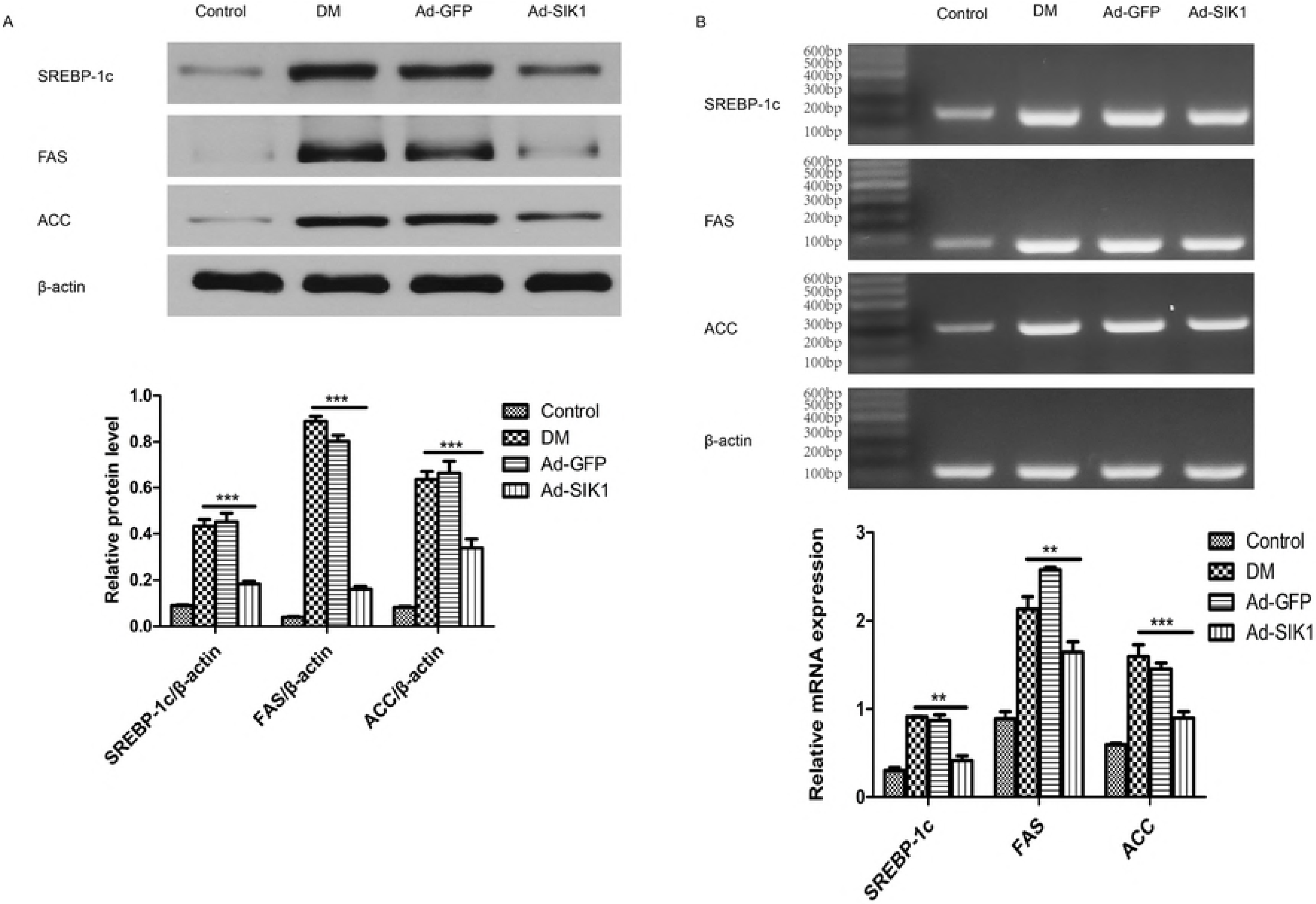
Effects on mRNA and protein expression of genes related to lipid metabolism in diabetic rats. (A) Protein levels of SREBP-1c, FAS and ACC in liver; (B) mRNA (relative fold change) levels of SREBP-1c, FAS and ACC in liver. Fold expression levels were measured relative to the expression of β-actin (internal control). Data are presented as the mean ± SD. Significant differences are indicated as * p<0.05, ** p<0.01, *** p<0.001. ns, not significant. n=6 in the control group; n=5 in the DM group; n=6 in the Ad-SIK1 group.

To investigate the underlying molecular mechanism of the hypolipidaemic effect of SIK1 overexpression on the diabetic rats, we measured the mRNA and protein of SREBP-1c, FAS and ACC in the liver. Notably, the mRNA expression of SREBP-1c, FAS and ACC in the liver of the DM group increased significantly compared to the control group (Fig 4B). However, Ad-SIK1 treatment significantly reduced the mRNA expression of SREBP-1c, FAS and ACC in the liver compared with the DM group, suggesting the mitigative role of SIK1 on fatty liver. Meanwhile, Western blot analysis revealed that SREBP-1c, FAS and ACC were markedly downregulated in the Ad-SIK1 group compared with the DM group (Fig 4A). Taken together, these findings indicate that the relieving effect of SIK1 overexpression on fatty liver was associated with a significant reduction in the expression of lipogenetic genes such as SREBP-1c, FAS and ACC.

## Discussion

Since the discovery of the SIK family, the roles of SIK isoforms (SIK1/2/3) in glucose and lipid metabolism have been extensively investigated [6–9, 12, 14, 17, 18]. However, the biological function of SIK1 remains poorly understood in HFD/STZ-induced T2DM rats. In this study, we found that the expression of hepatic SIK1 was markedly decreased in the HFD/STZ-induced T2DM rat model and that administration of Ad-SIK1 lowered fasting blood glucose and ameliorated fatty liver disease, suggesting that a reduction of SIK1 may contribute to the glucose and lipid metabolism disorder in diabetes. Metformin, a widely used hypoglycemic drug, which attenuated hyperglycaemia and NAFLD in HFD/STZ-induced diabetic rats [27], increased SIK1 expression levels in HepG2 cells cultured in high glucose conditions [18]. These findings indicate that SIK1 may be associated with the pathogenesis of T2DM and NAFLD.

This study established a model of T2DM rats treated with HFD plus low-dose STZ. The HFD/STZ-induced diabetic rats exhibited classic diabetic symptoms of polyuria, polydipsia and weight loss. The T2DM rats also showed high blood glucose, high TG and TC than normal control rats, which was consistent with previous report [26]. We further determined the effect of Ad-SIK1 on hepatic glucose metabolism in HFD/STZ-induced diabetic rats. Although the function of SIK1 in the HFD/STZ-induced diabetic rat model has not been reported, but RNAi-based knockdown strategies and genetic diabetic mouse models have revealed its role in the regulation of glucose metabolism. First, overexpression of SIK1 reduced fasting blood glucose and gluconeogenic gene expression in db/db mice, while Ad-SIK1 RNAi promoted them [9]. Second, SIK1 activity was reduced in the livers of db/db diabetic mice[12]. In the present study, the administration of Ad-SIK1 resulted in amelioration of hyperglycaemia in HFD/STZ-induced diabetic rats, suggesting that exogenous SIK1 might have a protective effect on T2DM. This is obviously the first report that suggests the potential of adenovirus-mediated SIK1 gene transfer in the management of hyperglycaemia in the HFD/STZ-induced T2DM rat model. More importantly, we for the first time demonstrated that SIK1 mRNA and protein expression were significantly reduced in the livers of HFD/STZ-induced diabetic rats, which was consistent with our previous in vitro studies [8, 18], suggesting that the expression of SIK1 is inhibited in diabetic states.

Phosphorylation at Thr182 by LKB1 is essential for switching on the SIK1 kinase activity [7, 28]. The Thr182 of SIK1 is phosphorylated by LKB1, resulting in conversion from inactive SIK1 to the active form [6]. Consistent with previous in vitro observations [8, 18], this in vivo study indicated that the level of Thr-182 phosphorylation, as well as the expression of SIK1 mRNA and protein, was downregulated in the livers of HFD/STZ-induced diabetic rats, suggesting that the SIK1 kinase activity may be suppressed in diabetic states. As expected, Ad-SIK1 treatment significantly elevated the level of pT182 SIK1 compared to the DM group. In addition, the intracellular distribution of SIK1 is closely associated with its functional activity. Ser-577 is a determinant of the intracellular distribution of SIK1. Moreover, inactive SIK1 as well as CREB-repressing active SIK1 are present as Ser577-dephosphorylated forms and are localized in the nucleus [3, 6, 8]. Phosphorylation of SIK1 at Ser577, which causes the nucleus export of SIK1, leads to a reduction of the transcriptional modulating activity of SIK1 [6]. On the basis of the ability of Ser 577 phosphorylation to decrease the transcriptional modulating activity of SIK1, we reasoned that phosphorylation level of Ser577 indicated the ability of SIK1 to suppress CREB. Thus, we examined the phosphorylation level of SIK1 at Ser577 in the livers of HFD/STZ-induced diabetic rats. In vivo, in the DM and Ad-GFP groups, the phosphorylation of SIK1 at Ser577 was elevated, whereas the expression of SIK1 was reduced, which were reversed by Ad-SIK1 administration, suggesting the possibility that SIK1 acts as a modulator of CREB-dependent transcription in the livers of HFD/STZ-induced diabetic rats.

The remarkable feature of T2DM is elevated fasting blood glucose. Dysregulated gluconeogenesis contributes to hyperglycaemia in diabetic rodents and humans [29, 30]. SIK1 was shown to inhibit CREB activity by phosphorylating CREB-specific coactivators, CRTC2, at Ser171 to suppress hepatic gluconeogenesis [9, 10]. Serine 171 is the primary phosphorylation site that mediates CRTC2 activity [9]. To confirm the importance of Ser 171 for inhibition of the gluconeogenic programme by SIK1, we evaluated the expression of hepatic SIK1, CRTC2 and pS171 CRTC2 in diabetic rats. Our results showed that the mRNA and protein expression of CRTC2 in the DM and Ad-GFP groups was significantly elevated, whereas pS171 CRTC2 was downregulated compared to the control group. Moreover, relative to control Ad-GFP diabetic rats, Ad-SIK1 administration decreased fasting blood glucose, increased pS171 CRTC2 and reduced CRTC2 and gluconeogenic genes, such as PEPCK and G6Pase. The changes in expression of SIK1, CRTC2, PEPCK and G6Pase were also confirmed by immunohistochemistry analysis. Interestingly, the CRTC2 nuclear accumulation observed in the Ad-SIK1 group was lower than that seen in the DM group, suggesting that Ad-SIK1 treatment might deter the translocation of CRTC2 into the cell nucleus, thus reducing the transcription of gluconeogenic genes and hepatic glucose output. Consequently, we observed lower blood glucose levels in the Ad-SIK1 group than in the DM group. Taken together, these findings suggest that recombinant SIK1 directly inhibited the hepatic gluconeogenic program in the HFD/STZ-induced diabetic rats by the SIK1/CRTC2 pathway.

Lipid metabolic disorder is one of the most common pathophysiological changes in T2DM. Liver plays a vital role in the regulation of systemic lipid metabolism. As a key regulator of hepatic lipogenesis, SREBP-1c was suggested to be involved in the development of NAFLD by contributing to the onset of fatty liver phenotypes [17]. SIK1 regulates hepatic lipogenesis by modulating SREBP-1c activity [17]. To evaluate the effect of adenovirus-mediated SIK1 overexpression on lipogenic gene expression in the livers of HFD/STZ-induced T2DM rats, we transduced diabetic rats with Ad-SIK1 adenovirus or Ad-GFP control viruses. In this study, the HFD/STZ-induced diabetic rats showed characteristics of NAFLD, including elevation of hepatic enzyme levels, significantly increased relative liver weights (liver index), hyperlipidaemia and histological changes such as steatosis and hepatocyte injury. In concert with the histological and immunohistochemistry analysis, Ad-SIK1 administration reduced serum TC and TG levels, and decreases the elevated hepatic mRNA and protein levels of SREBP-1c, FAS and ACC caused by HFD/STZ-induced T2DM, suggesting that overexpression of SIK1 could suppress hepatic lipogenesis by downregulating SREBP-1c and its downstream gene expression. This effect was consistent with previous reports [12, 17].

In summary, the present study demonstrates that SIK1 mRNA and protein expression are significantly reduced in the livers of HFD/STZ-induced diabetic rats. Overexpression of SIK1 ameliorates hyperglycaemia and fatty liver by suppressing hepatic gluconeogenesis and lipogenesis in HFD/STZ-induced T2DM rats. This protective effect of SIK1 may be derived from its interference with the SIK1/CRTC2 and SIK1/SREBP-1c pathways. Up-regulating hepatic SIK1 expression may represent an attractive means for the treatment of T2DM and NAFLD. Figure 6 illustrates the possible mechanisms of SIK1 in attenuating T2DM with NAFLD.

**Figure 6.**
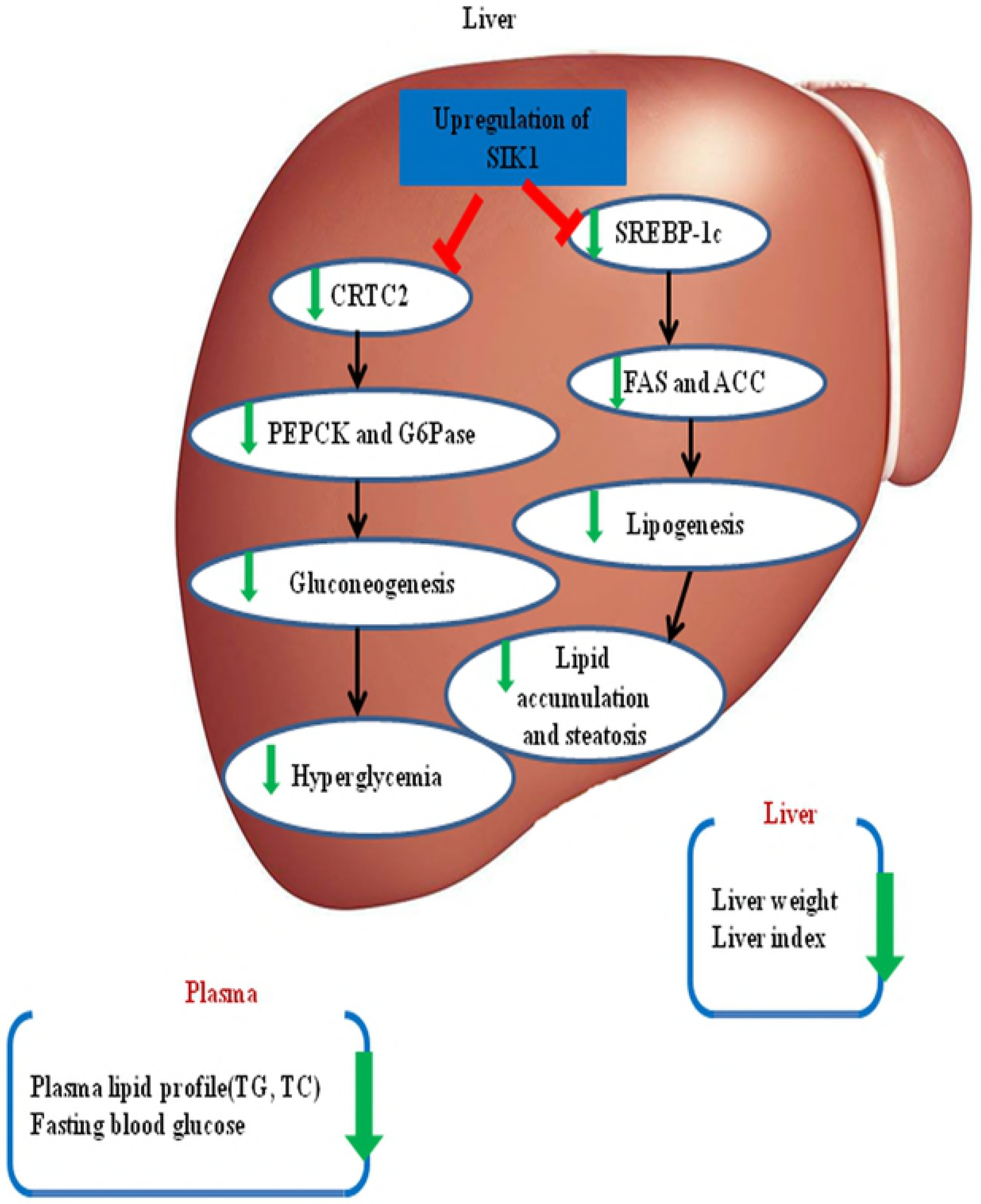
Proposed mechanisms of the hypoglycemic and hypolipidemic effect of SIK1. Schematic representation of the role of SIK1 in amelioration of T2DM with NAFLD. Overexpression of SIK1 can contribute to preventing hyperglycemia and hepatic lipid accumulation through suppressing gluconeogenesis and lipogenesis with a decrease in the expressions of CRTC2, PEPCK, G6Pase, SREBP-1c, FAS and ACC in liver, thus reducing fasting blood glucose, serum TC, serum TG, hepatic steatosis and liver weight. SIK1, salt-induced kinase 1; CRTC2, CREB-regulated transcription co-activator 2; PEPCK, phosphoenolpyruvate carboxykinase; G6Pase, glucose-6-phosphatase; SREBP-1c, sterol regulatory element binding-protein-1c; ACC, acetyl-CoA carboxylase; FAS, fatty acid synthase; TC, total cholesterol; TG triglycerides.

## Supporting Information

**S1 Appendix.** Method of construction of recombinant adenovirus vectors.

**S1 Table.** List of primer sequences for RT-PCR.

**S2 Table.** Changes in body weight, liver weight, FBG, TG and TC.

## Author Contributions

Conceptualization, XiuYing Wen; Data curation, DaoFei Song and Chang Wang; Formal analysis, DaoFei Song and Lei Yin; Funding acquisition, XiuYing Wen; Investigation, DaoFei Song; Methodology, XiuYing Wen; Project administration, DaoFei Song; Supervision, XiuYing Wen; Validation, DaoFei Song; Writing – original draft, DaoFei Song; Writing – review & editing, XiuYing Wen.

## Abbreviations

ACC: acetyl-CoA carboxylase
ACTH: adrenocorticotropic hormone
Ad-GFP: adenovirus-green fluorescent protein
Ad-SIK1: adenovirus-Salt induced kinase 1
AMPK: AMP-activated protein kinase
CREB: cAMP response element binding protein
CRTC2: CREB-regulated transcription co-activator 2
DM: diabetes mellitus
FAS: fatty acid synthase
G6Pase: glucose-6-phosphatase
HFD: high-fat diet
LKB1: serine/threonine kinase 11
NAFLD: nonalcoholic fatty liver disease
OD: optical density
PGC-1α: peroxisome proliferator-activated receptor gamma coactivator 1-alpha
PKA: protein kinase A
PEPCK: phosphoenolpyruvate carboxykinase
PFU: plaque forming units
RT-PCR: reverse transcription-polymerase chain reaction
SIK1: Salt-induced kinase 1
STZ: streptozotocin
SREBP-1c: regulatory element binding-protein-1c
T2DM: type 2 diabetes mellitus
TC: total cholesterol
TG: triglycerides

